# Climatic niche conservatism in non-native plants depends on introduction history and biogeographic context

**DOI:** 10.1101/2025.01.28.635214

**Authors:** Anna Rönnfeldt, Valén Holle, Katrin Schifferle, Laure Gallien, Tiffany Knight, Patrick Weigelt, Dylan Craven, Juliano Sarmento Cabral, Damaris Zurell

## Abstract

Many tools informing preventive invasion management build on the assumption that introduced species will conserve their climatic niches outside their native ranges. Previous research testing the validity of this assumption found contradictory results regarding niche conservatism vs. niche switching for non-native species. An open question is in how far these contradictions reflect context dependency, yet only few studies compared the niche dynamics of species introduced to multiple regions. Here, we used an ordination-based approach to quantify the climatic niche changes (stability, unfilling, expansion) of 316 plant species introduced to eight different regions across the world, including the Pacific region with extreme isolation between island groups. We then performed multiple phylogenetic regressions to assess how the regional context and species’ characteristics affect niche dynamics. Niche conservatism varied across regions, even within species. While niche expansion into previously unoccupied climates was generally low, niche unfilling varied strongly between regions. Generally, region-specific introduction history and species’ biogeographic attributes were more important for explaining niche changes than ecological traits. Niche expansion was consistently higher for species with small native range sizes, and niche stability increased. In contrast, niche unfilling decreased with time since introduction which could suggest that the lack of niche conservatism observed in many regions might be transient and potentially related to dispersal limitations. Overall, our results shed light on the context dependency of climatic niche changes when species are introduced to new regions, highlighting that the species and region-specific context should be accounted for when assessing the potential for niche changes.

**Significance Statement:** As non-native species are introduced to new regions by humans, they may occupy the same climatic conditions as in their native ranges, leave parts of their native niche unfilled or expanding into previously unoccupied conditions. Knowing to which extent these niche dynamics generally occur is essential for understanding and managing biological invasions. Here, we evaluate regional differences in the climatic niche dynamics of non-native plants that were introduced to multiple regions across the world. We found marked variation across regions, influenced by factors such as the time that has passed since species were introduced, or biogeographic attributes of both the native and non-native ranges. These findings indicate that a lack of apparent niche changes is likely a temporary phenomenon.

## Introduction

Invasive species threaten biodiversity and human well-being worldwide (1, 2). Given the expansion of global trade and the limited funding for conservation, preventing the introduction and establishment of non-native species is the most cost-effective option to manage invasive species and to reduce their negative impacts (3–5). Targeted prevention can be facilitated by model-based risk assessments that help guide monitoring and early detection. Correlative species distribution models (SDMs, 6) are particularly prominent in global change research to infer non-native species’ realized climatic niches from empirical data and to predict potential climatic suitability in space and time. Importantly, these models rely on the assumption that species occupy the same climatic space in their region of origin as in their invaded regions: so-called niche conservatism (7). Thus, our ability to predict the potential establishment of non-native species is entangled with the question of whether species conserve their niche when introduced elsewhere.

Several processes can influence the degree to which species conserve their niche during an invasion, which would then be reflected through different metrics in a niche comparison (Fig. 1). If species are at equilibrium and conserve their niche, meaning they occupy the same climatic conditions in the introduced ranges as in their native ranges, they should show high niche stability. An ongoing colonization process in a new region and dispersal barriers that hinder species from reaching suitable climatic conditions, would cause species to occupy a smaller subset of their climate niche in their introduced range, known as niche unfilling (8, 9). Niche expansion into previously unoccupied parts of the climatic niche space can occur due to changes in biotic interactions, e.g., through competitive or enemy release (10, 11), the absence of dispersal barriers present in the native range, or rapid adaptation occurring after the introduction to a new region (12). Niche stability, expansion and unfilling relate to the analogue niche space (climatic conditions present in both ranges; Fig. 1)(13). Other niche metrics related to non-analogue climates (niche abandonment and niche pioneering) are difficult to compare between species and regions. To date, evidence for or against niche conservatism in non-native ranges is controversial (e.g., 9, 14–17), but methodological differences across studies make direct comparisons difficult (8). A promising way forward is to assess climatic niche changes across species that have been introduced to multiple regions, which would serve as a natural experiment to identify potential generalities in the niche dynamics of non-native species.

**Figure 1.**
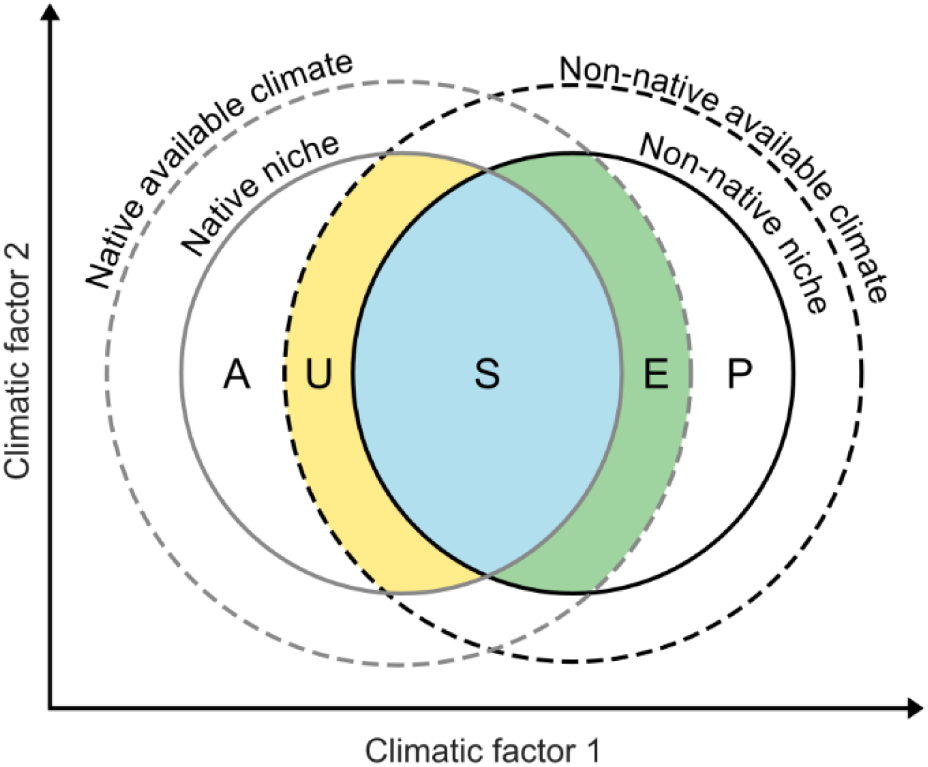
Conceptual overview of subcomponents of niche differences between the native and non-native range. The solid lines indicate the native (grey) and non-native (black) climatic niches and the dashed lines the available climatic conditions in the native and non-native range. In analogue climatic conditions, niche stability (S, blue) describes the parts of the climatic niches occupied in both ranges, unfilling (U, yellow) indicates native niche space that is not part of the non-native niche, and expansion (E, green) refers to non-native niche space that is not part of the native niche. Under non-analogue conditions, abandonment (A) indicates environmental conditions within the native niche that are not available in the non-native range, and pioneering (P) describes the part of the non-native niche that has no analogue in the native range. Figure adapted from Fig. 1 in Guisan et al. (13).

Multiple factors have the potential to affect the niche dynamics a species will display in a region: the time that has passed since the initial introduction (residence time), the species’ functional traits, and species-specific biogeographic attributes related to the native and non-native ranges (for simplicity from here on referred to as biogeographic attributes). Niche unfilling, for example, has been found to be lower for non-native vertebrate species with comparably longer residence times (18). Higher dispersal ability, reflected for example by traits associated with long-distance dispersal, could aid non-native range expansion (19) and result in relatively lower niche unfilling compared to dispersal-limited species. According to Early and Sax (20), higher niche expansion might be found in species with small native ranges and narrow climatic niches because of release from non-climatic niche and range constraints, such as biotic interactions (e.g., fungal pathogens, 21). We also hypothesize latitudinal distance between the native and non-native ranges to affect the niche dynamics. A longer latitudinal distance between the regions could be indicative of divergent biogeographic history, environmental and biotic filtering and, consequently, lower niche stability in the non-native range. While ecological traits are intrinsic to species, some of their biogeographic attributes and introduction history may vary across non-native regions, increasing the likelihood of regional differences in the niche dynamics and the level of niche conservatism (Table 1).

**Table 1.**
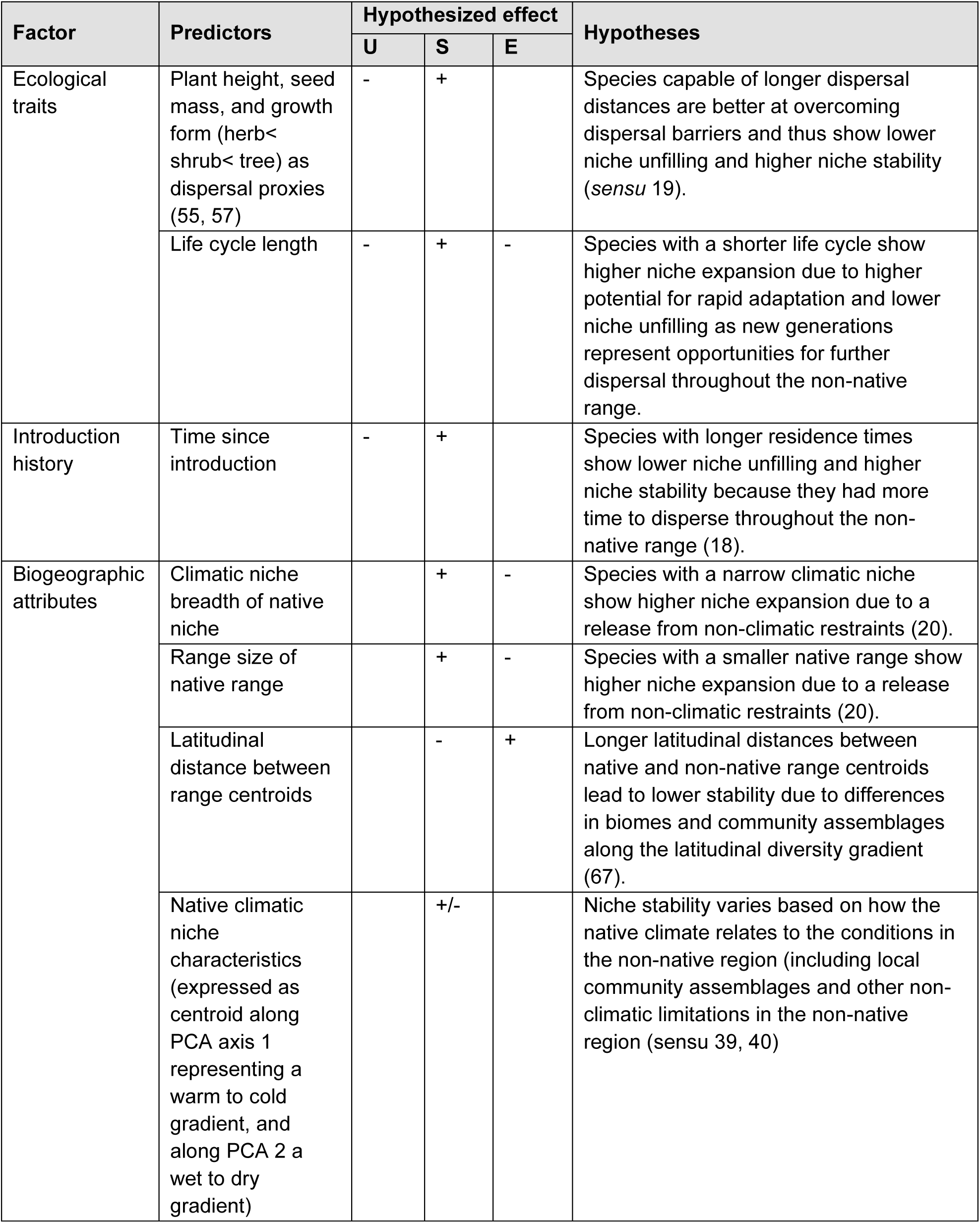
Overview of the hypothesized factors that might affect niche unfilling (U), stability (S) or expansion (E) of non-native plants upon their introduction to a new range. The hypothesized effects of the predictors on each niche change metric can be positive (+), negative. (**-**)**, mixed (+/-) or non-existent.**

| Factor | Predictors | Hypothesized effect |  |  | Hypotheses |
| --- | --- | --- | --- | --- | --- |
|  |  | U | S | E |  |
| Ecological traits | Plant height, seed mass, and growth form (herb< shrub< tree) as dispersal proxies (55, 57) | - | + |  | Species capable of longer dispersal distances are better at overcoming dispersal barriers and thus show lower niche unfilling and higher niche stability (*sensu* 19). |
|  | Life cycle length | - | + | - | Species with a shorter life cycle show higher niche expansion due to higher potential for rapid adaptation and lower niche unfilling as new generations represent opportunities for further dispersal throughout the non-native range. |
| Introduction history | Time since introduction | - | + |  | Species with longer residence times show lower niche unfilling and higher niche stability because they had more time to disperse throughout the non-native range (18). |
| Biogeographic attributes | Climatic niche breadth of native niche |  | + | - | Species with a narrow climatic niche show higher niche expansion due to a release from non-climatic restraints (20). |
|  | Range size of native range |  | + | - | Species with a smaller native range show higher niche expansion due to a release from non-climatic restraints (20). |
|  | Latitudinal distance between range centroids |  | - | + | Longer latitudinal distances between native and non-native range centroids lead to lower stability due to differences in biomes and community assemblages along the latitudinal diversity gradient (67). |
|  | Native climatic niche characteristics (expressed as centroid along PCA axis 1 representing a warm to cold gradient, and along PCA 2 a wet to dry gradient) |  | +/- |  | Niche stability varies based on how the native climate relates to the conditions in the non-native region (including local community assemblages and other non-climatic limitations in the non-native region (*sensu* 39, 40) |

In addition to species-specific factors, the regional context may affect niche dynamics. Species introductions to islands, for example, could result in more pronounced differences in the niche dynamics compared to mainland regions. The geographic isolation of islands could lead to dispersal limitation and higher niche unfilling, or the depauperate species communities of islands could result in higher niche expansion due to a higher chance of biotic constraints being lifted (22). One such island system is the Pacific Island region, which includes thousands of islands across the Pacific Ocean. The region is known for its high vulnerability to biological invasions (23) and a steep increase in invasive plant species per unit area (24). Many of the non-native plants that have been introduced to the Pacific Islands have also been introduced elsewhere in the world (25). This marks the region as an ideal natural laboratory to quantify regional differences in niche dynamics.

In this study, we pursue two objectives. First, to quantify and compare the climatic niche changes between native and non-native ranges of vascular plant species that have been introduced to multiple regions of the world, with a particular focus on potential differences between the Pacific Islands compared to other regions. Second, to test how ecological traits, biogeographic attributes, and region-specific introduction history affect the magnitude of the niche dynamics. We considered vascular plant species included in the PaciFLora data set (26), which lists known naturalized occurrences of non-native species on the Pacific Islands. We matched georeferenced occurrence data of 316 plant species with a biogeographic status (native or non-native) across eight study regions. We then quantified niche change using an ordination-based approach. Finally, we performed phylogenetic multiple regression to assess the relationship between the niche change metrics and ecological and biogeographic traits, as well as species and region-specific introduction history (Tab. 1).

## Results

### Niche conservatism

We followed the methodology of Broennimann et al. (27) to quantify native and non-native niche spaces along the first two axes of a principal component analysis (PCA) over 19 bioclimatic variables, in order to assess niche similarity and to quantify niche change metrics between pairs of native and non-native niches. We compared 1566 niche pairs, capturing the introductions of 316 species across up to eight botanical continents (28): the Pacific Islands, Africa, temperate Asia, tropical Asia, Australasia, Europe, North America and South America. The native ranges of these species often covered multiple climatic zones (classified based on the main Köppen climate zones, 34) and regions (SI appendix). The most common climate zones found in the native ranges were the tropical, temperate, and a combination of tropical and temperate zones, for around 30 % of the species respectively. There was an even flow of species between the different native climate zones and the non-native regions, except for fewer temperate species having been introduced to tropical Asia and fewer introductions of tropical species to Europe (Fig. S2).

We observed significant niche conservatism for 45.5 % of the introductions and found no indication of niche switching (the opposite of niche conservatism) according to the null-model-based similarity test (Fig. S3). For most niche pairs, however, the similarity test results were not statistically significant, meaning that these species neither conserve nor switch their climatic niche but showed an intermediate degree of niche changes. Notably, most species showed inconsistent patterns of niche conservatism across regions, with significant niche conservatism in some but not all non-native regions (269 out of 316 species, Fig. 2). Only 20 species consistently conserved their climatic niche across all regions they were introduced to, while 47 showed no niche conservatism in any region (Fig. 2). The lowest proportion of species with significant niche conservatism was found in the Pacific region (Fig. 2).

**Figure 2.**
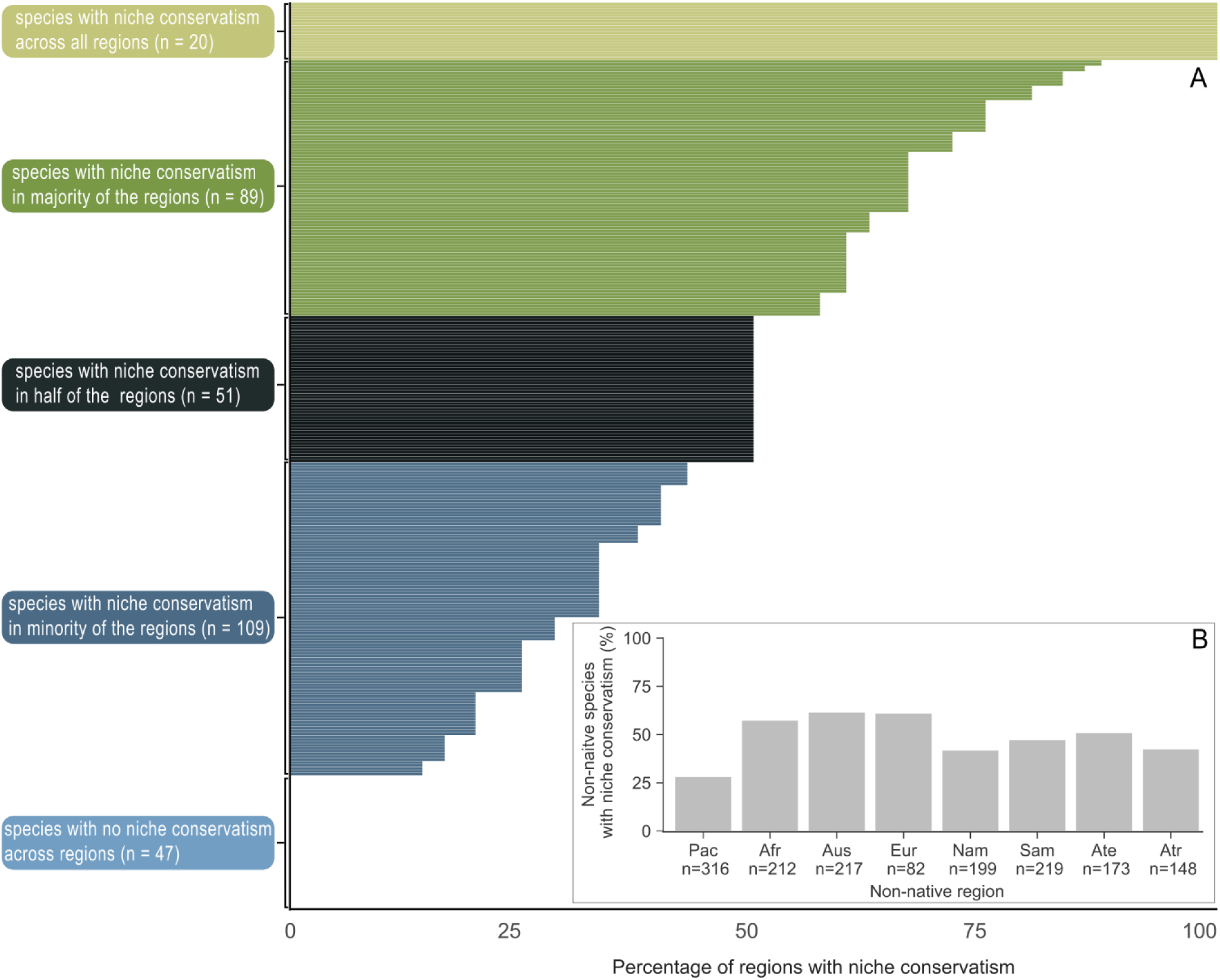
Niche conservatism across regions. Similarity tests were used to test for significant niche conservatism in non-native vascular plants introduced to different regions. In panel (A), each horizontal line represents one species and the percentage of regions in which that species showed significant niche conservatism based on null models (n = 1200 iterations). Colours indicate whether species conserved their niche in all (light green), the majority (≥ 50%, dark green), half (black), the minority (≤ 50%, dark blue) or none of the regions in which they have been introduced. Panel (B) shows the percentage of species that conserved their niche for each non-native region respectively (Pac = Pacific Islands, Afr = Africa, Aus = Australasia, Eur = Europe, Nam = North America, Sam = South America, Ate = temperate Asia, Atr = tropical Asia). Note that no species showed statistically significant niche switching (Fig. S3).

### Niche changes

We focused our analyses on the analogue niche change metrics stability, unfilling and expansion, with additional results for niche abandonment and pioneering provided in the SI appendix. Niche stability was high across species and regions (median 78.4 %). Many species exhibited niche unfilling (median 17.6 %), while niche expansion was relatively low overall (median 0.78 %). ANOVA results indicated significantly higher niche unfilling for plants introduced to the Pacific Islands compared to all other regions, except North America (Fig. 3, Tab S1). Consequently, niche stability in species introduced to the Pacific Islands was often lower compared to the other regions. Niche expansion was low overall, with the highest median niche expansion reaching 3.4 % in tropical Asia. However, we observed several outliers in each region, with some species showing niche expansion exceeding 50 % (SI appendix).

**Figure 3.**
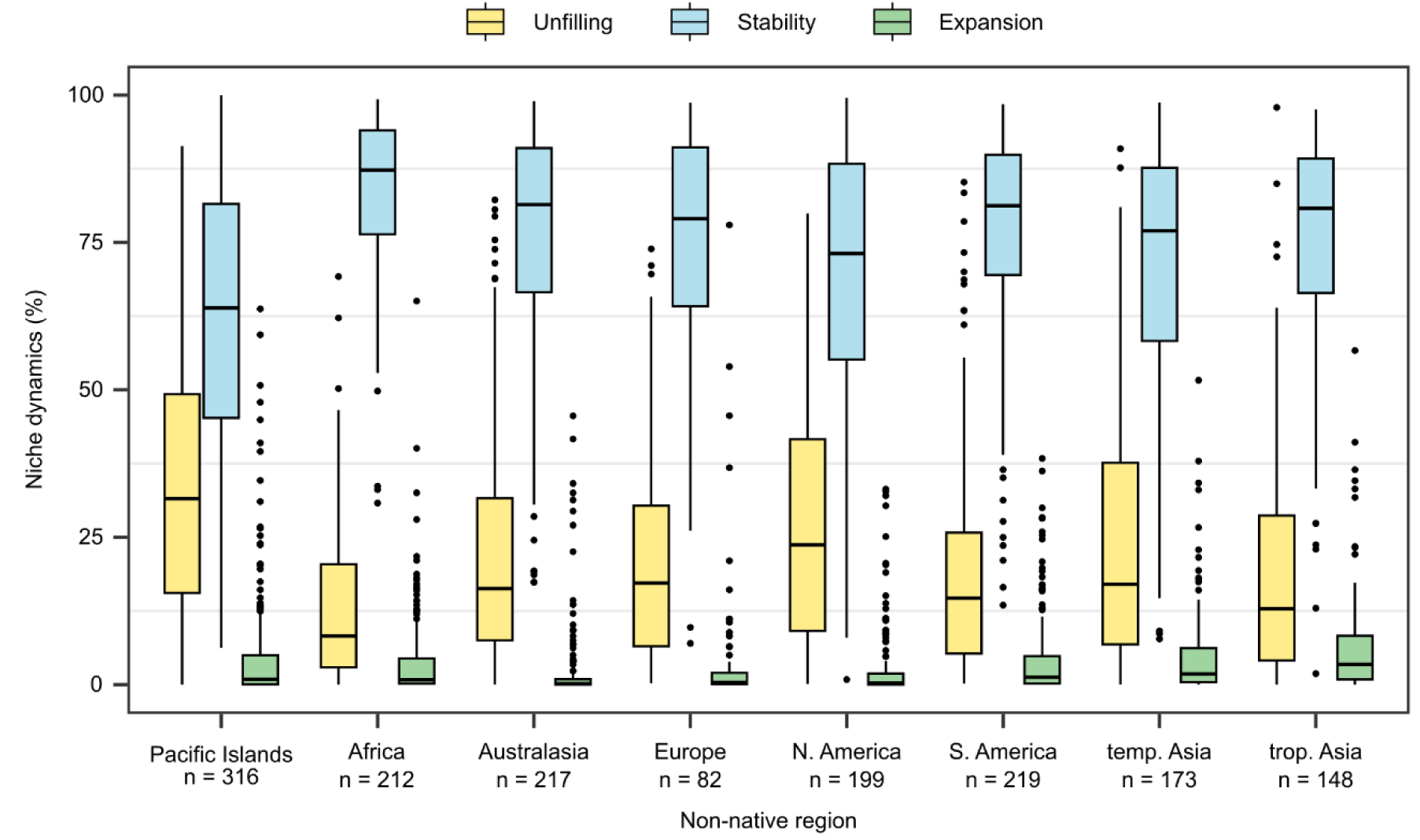
Climatic niche dynamics in non-native plants across regions. The sample size in each region indicates the number of species that have been introduced to the respective region.

### Trait analyses

We used multiple phylogenetic regression to assess the effect of ecological traits, biogeographic attributes and residence times on the region-specific niche change metrics while controlling for shared evolutionary history between species. Due to data gaps, these analyses were restricted to 165 of the 316 plant species, covering 770 of the 1566 niche pairs. Species from the different climate zones had overall similar mean trait values, except for a wider mean niche breadth in temperate species and a higher mean seed mass in tropical species (Fig. S5). Separate models were fitted for each niche change metric and region. Trait models explained 7-52 % of the variance in niche unfilling (Fig. 4A), 17-56 % in niche stability (Fig. 4B), and 17-55 % in niche expansion (Fig. 4C). The relative importance of different traits varied considerably across regions and considered niche change metrics. Biogeographic attributes and species residence time were overall more important than ecological traits for explaining niche changes (Fig. 4). A few general patterns emerged: niche unfilling consistently decreased and niche stability increased for species with longer residence times (Fig. 4A-B). Also, niche expansion consistently decreased for species with larger native range sizes and increased with longer distances between latitudinal range centroids (Fig. 4C). These patterns also emerged in the univariate models, indicating that they are robust against missing species (Fig. S8).

**Figure 4.**
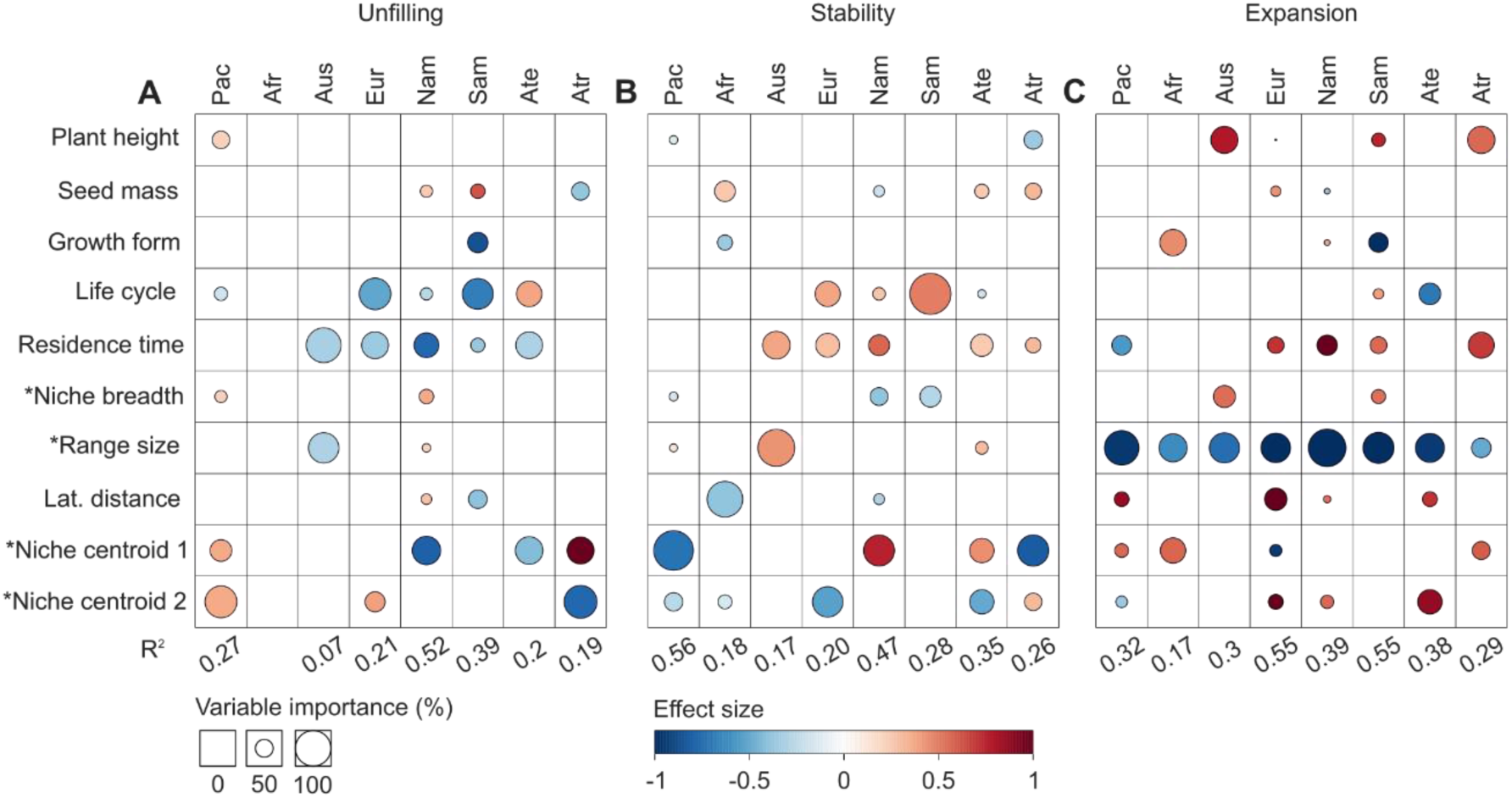
Trait effects on climatic niche changes in non-native plants. For each non-native region, we fitted regression models with AIC-based stepwise selection for (A) niche unfilling, (B) niche stability and (C) niche expansion. Each column represents the final model for each region, with the total explained variance (R^2^) shown below the columns. Circle sizes indicate the variable importance (%) of single traits within the multiple regression models, and the effect size shows whether the respective niche change metric increases (red) or decreases (blue) as the trait values increase. Traits labelled with an asterisk refer to biogeographic traits estimated for the native range or native niche of the species. Niche centroids refer to the relative position along climatic gradients 1 (from warm to cold) and 2 (from wet to dry). The sample size varied between regions: Africa (Afr, n = 124), temperate Asia (Ate, n = 95), tropical Asia (Atr, n = 78), Australasia (Aus, n = 124), Europe (Eur, n = 56), North America (Nam, n = 110), Pacific Islands (Pac, n = 143), South America (Sam, n = 41).

Among the biogeographic attributes, niche centroids were generally most important for explaining niche unfilling and niche stability, but the direction of these effects was variable (Fig. 4). For example, species native to colder climates (niche centroid 1) showed higher niche stability and less niche unfilling when introduced to temperate Asia and North America, but lower niche stability and stronger unfilling when introduced to tropical Asia and the Pacific Islands. Species native to dry climatic conditions (niche centroid 2) showed lower niche stability and more niche unfilling when introduced to Europe and the Pacific Islands, and higher niche stability and lower niche unfilling when introduced to tropical Asia. While native range size negatively affected niche expansion in all regions (Figs. S5-S6), it was rarely selected in the final trait models for niche stability and niche unfilling (Fig. 4). Also, the effects of the native climatic niche breadth and latitudinal distance between native and non-native regions were minor. If selected in the final models, however, climatic niche breadth had a negative effect on niche stability and a positive effect on niche expansion and niche unfilling (Fig. 4). For the ecological traits, no generalities emerged. Where life cycle was included in the final models, niche unfilling tended to decrease and niche stability to increase for longer-lived species, except in temperate Asia where the opposite was observed. Niche expansion, surprisingly, increased with plant height in a few regions (Fig. 4).

## Discussion

Our niche comparison quantifying the climatic niche changes of non-native plants introduced to different world regions revealed that climatic niche conservatism is highly context-dependent and largely depends on residency time. The magnitude of niche changes strongly varied across regions. Niche stability was generally high, although substantial niche unfilling occurred while niche expansion was consistently low. Using null models, we found that the same species may show significant niche conservatism in some regions but not in others. We further found that biogeographic attributes and time since introduction affected niche changes more strongly than ecological traits linked to dispersal and life cycle. This suggests that climatic niche conservatism depends, at least to some extent, on the climatic preferences of the introduced species and that climatic niche shifts may be primarily transient.

We found strong variation in climatic niche changes across regions, indicating that regional inconsistencies in niche changes are common among plants. This corroborates previous findings from multi-region niche comparisons of a few selected plant species (30, 31). Niche unfilling was prevalent across all regions, which could be due to geographic barriers hindering dispersal or biotic constraints. It could also indicate transient dynamics resulting from ongoing dispersal processes after the species’ initial introduction (8, 9). As hypothesized, we found that niche unfilling was higher for introductions to the Pacific than to other regions, likely due to the barrier effects of the ocean. It suggests that many of the species introduced to the Pacific Islands have not yet reached their climatic equilibrium and have unrealized colonization potential (32). It is thus likely that human-mediated transport, which often facilitates inter-island dispersal in the Pacific region (25) will eventually enable some of these species to fill their potential range, further increasing the invasion pressure on this region.

Niche expansion was generally low indicating that competitive release or niche adaptation is not prevalent across species and regions. In the case of the Pacific Islands, where we expected to see more niche expansion because of the depauperate species communities of islands (22), the geographic fragmentation of the region could have further contributed to the species not being able to reach and thus expand into otherwise available niche spaces. To understand the specific mechanisms behind the low expansion values a more small-scale study would be more suitable, focusing on factors such as regional community assemblages, the total climatic expansion potential (33), or anthropogenic transport networks facilitating inter-island dispersal (25). When comparing the overall low niche expansion with previous studies that quantified niche differences across several continents, our results stand in stark contrast to results by Atwater et al. (14) who found niche expansion to be common in introduced plants. In contrast, our results align with Petitpierre et al. (9) who also found lower niche expansion compared to niche unfilling. Low niche expansion would be a promising sign for the reliability of model-based risk assessments that typically rely on the assumption that the realized niche is adequately captured by data (7, 34), especially for non-native species where no georeferenced data on previous introductions to other regions are available. However, it is important to note that comparably little niche expansion in climatic space could still result in a high species prevalence in geographic space if those climates are common (35). This could lead to an underprediction of the establishment potential of non-native species (36, 37) and thus to uncertainty in management planning and decision-making. While over-predictions rather than under-predictions are more desirable for risk assessments, pronounced niche unfilling in the geographic space can likewise affect management if the costs of preventive measures across larger areas would drastically outweigh the eventual negative impacts of a species should it ever fill its entire niche in these areas (38).

Despite consistent evidence of climatic niche changes across species and regions, we did not detect significant niche switching. Still, we confirmed significant niche conservatism in less than half of the considered introductions, and we found many cases where species showed significant niche conservatism only in one or few (but not all) regions. This implies that the contradictory results found in the existing literature on niche conservatism in introduced species might not only be a result of methodological differences and the focus on different species (8, 13) but also due to differences across focal regions. Although recent reviews indicate that niche conservatism is the norm when species are introduced to a new range (8), we did not find direct support for this in our data, as niche conservatism was not a general trend for the species introduced to multiple regions. We do, however, suggest that a lack of niche conservatism is likely temporary for many of these introductions as our trait analysis revealed a strong connection between niche unfilling and a species’ residence time in a region. This corroborates previous findings by Strubbe et al. (18) and emphasizes the transient nature of niche unfilling. Interestingly, although residence time also had a negative effect on introductions to the Pacific Islands, it was not selected in the final models, indicating that compared to other regions residence time had less effect in the Pacific. This is likely due to the geographic isolation between islands drastically limiting natural dispersal across the region. For example, *Ricinus communis* (L.), one of the species with the longest residence times in our data set, only showed 1.6 % of niche unfilling in North America, 201 years after its initial introduction. Only three years prior, *R. communis* was first recorded on one of the Pacific Islands, yet, unfilling still contributes 43.4 % to the analogue niche space of the region. Overall, the high prevalence of niche unfilling and associated dispersal limitations and the strong role of time since introduction for explaining these phenomena further corroborates that niche conservatism is a matter of time and that most species could eventually tend towards climatic niche conservatism. Likely exceptions to this are introductions to archipelagic systems unless anthropogenic activities bridge the inter-island dispersal limitations.

Our trait analyses additionally revealed that niche conservatism is highly context-dependent. Specifically, the native climatic niche position had a consistently strong effect on niche stability and niche unfilling. Thus, niche stability or conservatism is to some degree determined by the overlap between a species’ climatic preference and the main climatic conditions found in the non-native regions. Here we focused on the analogue niche space, so this does not refer to the presence or absence of climatic conditions. Rather, this connection likely reflects the importance of factors such as the occurrence of these analogue climatic conditions in the geographic space (common vs. marginal) and resulting non-climatic limitations (e.g., biotic interactions (39, 40) or land use intensity (41)) for the establishment and thus niche stability of an introduced species. This context dependency also emphasizes that observed niche changes in one region cannot be easily generalized to other regions. Another important biogeographic attribute affecting niche change was range size, which was often the most important variable per region. Species with smaller native range sizes, such as *Opuntia ficus-indica* (L.) native to Mesoamerica, showed comparably high niche expansion values in all eight study regions. This result aligns well with other studies that found native range size and niche expansion to be negatively correlated (20, 42). It indicates that the native ranges of these species were not constrained by climatic conditions (43) and that it is more likely that it was the release from non-climatic constraints (biotic or geographic) that enabled these species to expand their climatic niche in the non-native regions (10), rather than local adaptations upon introduction.

The effects of other traits on niche expansion and niche unfilling were less pronounced and varied more strongly across regions. Similar to range size, native niche breadth has previously been found to be negatively correlated with niche expansion (20). In our analyses, however, niche breadth was only relevant for explaining niche expansion in two regions and the effect direction was contrary to previous reports, with niche expansion increasing with climatic niche breadth. Also, we found little effect of ecological traits. We expected to observe less niche unfilling for species capable of longer dispersal distances and more niche expansion for species with shorter life cycles, but neither of these hypotheses was unequivocally corroborated by our results. This aligns with previous research by Early and Sax (20), who also found little effect of dispersal ability on niche unfilling. Overall, our trait analysis revealed that there are only a few general factors that explain climatic niche changes in non-native plants across regions. In particular, biogeographic attributes proved most important for explaining niche conservatism. Previous studies have shown that such biogeographic attributes are also critical determinants of the establishment success of introduced species (44, 45).

Multi-region comparisons of niche dynamics in non-native species are a major challenge and our analyses relied on a huge effort of consolidating heterogenous data sources for obtaining native and non-native occurrence information. We provide evidence that climatic niche conservatism may be a common endpoint of introductions into non-native regions and many species show more niche unfilling, which is indicative of dispersal limitations rather than niche expansion. Although outliers, some species can reach high levels of climatic niche expansion that appear to be related to changes in the realized climatic niche, for example, due to competitive release. Still, local adaptations, particularly on islands, cannot be completely discarded. Importantly, our results suggest that earlier contradictory findings of niche conservatism vs. niche switching could largely be related to context dependency and introduction history. Future research should thus more strongly focus on studying temporal aspects of climatic disequilibria in non-native species.

## Materials and Methods

### Study region

We focused on non-native plant introductions to eight distinct regions: the Pacific Islands, Africa, Australasia, Europe, North America, South America, temperate Asia, and tropical Asia (Fig. S1). Except for the Pacific Islands, these regions correspond to level 1 of the world geographic scheme of recording plant distributions (28). The spatial delimitation of the Pacific Islands was based on Wohlwend et al. (26) covering 50 island groups located in the Pacific Ocean between 40°N and 40°S.

### Species occurrence data

The initial species list consisted of the 3962 vascular plant species listed by the PaciFLora data set (26). PaciFLora builds on PIER (Pacific Island Ecosystems at Risk, http://hear.its.hawaii.edu/ Pier/) and GloNAF (Global Naturalised Alien Flora, 46) and lists vascular plant species with known naturalized occurrences on at least one of the islands in the Pacific region. We downloaded all available occurrence data for these species from GBIF (Global Biodiversity Information Facility, https://www.gbif.org/) and BIEN (Botanical Information and Ecology Network, https://bien.nceas.ucsb.edu/bien/) in June 2023, using the R packages *rgbif* (47) and *BIEN* (48). After merging these data at a 1 km resolution, we cleaned them by removing duplicates and occurrences with erroneous time stamps or coordinates, using the R package *CoordinateCleaner* (49). We then sampled background data within a 200 km terrestrial spatial buffer around the presence points to represent biogeographically accessible areas, aiming for a ratio of ten times more background data points than presence points, if possible (50). To reduce spatial autocorrelation, all presence and background data points were spatially thinned to a minimum distance of 3 km between points.

### Status assignment

Three data sources were incorporated to determine whether an occurrence point would be considered a native or non-native occurrence of the species in the respective regions. The first source was the WCVP (World Checklist of Vascular Plants, 51), which provides status information at a spatial resolution corresponding to level 3 of the world geographical scheme for recording plant distributions (28). To supplement the information provided by WCVP, we further included data from GIFT (Global Inventory of Floras and Traits, 52) and GloNAF (46). As the three data sources used different labels such as naturalized, alien, invasive, and non-native, we opted to only distinguish between *native* and *non-native* while harmonizing the information provided by the three sources. Conflicting status information was resolved (where possible) by using the status provided by the source that refers to a smaller spatial scale (e.g., a regional checklist rather than an entire country). Occurrences where conflicts between data sources could not be resolved or for which no matching status information was available were excluded.

### Climatic data

We used all 19 bioclimatic variables from CHELSA V2 (53, 54) at a 1 km resolution. These variables describe means, extremes, and seasonality of temperature and precipitation. The CHELSA V2 data are based on a mechanistic terrain-based downscaling approach and thus facilitate improved regional climate space estimates.

### Species trait data, residence time and biogeographic attributes

For subsequent trait analyses, we obtained data on ecological traits, biogeographic attributes, and the region-specific introduction history for each species. First, we considered four ecological traits: life cycle as a proxy for adaptive potential, and plant height, growth form and seed mass as proxies for dispersal ability (55). We gathered these data from the GIFT database, version 2.2, accessed via the R package *GIFT* (56). Categorical trait values were transformed to an ordinal scale. Life cycle ranged from short to long life cycles: annual < biennial < perennial. Growth forms ranged from short to long-distance dispersal potential (55, 57): herb < shrub < tree. Mean plant height (m) and seed mass (g) were continuous.

Second, we quantified several biogeographic attributes for each species: native niche breadth and centroid, native range size, and the region-specific latitudinal distance between native and non-native range centroids. We quantified native niche breadth using the Shannon Index of the species’ occurrence density in a two-dimensional climatic niche space (58, 59) using global climate as background data to allow inter-species comparisons. Occurrence density was corrected for climatic availability. Also, we extracted native niche centroids along the first two PCA axes (again estimated on global climate). The first axis described a gradient from warm to cold conditions (niche centroid 1) and the second axis a wet-to-dry gradient (niche centroid 2). Native range size was defined as the total area of the WCVP level 3 polygons and/or the GIFT polygons, with which native occurrence points were matched during the status assignment. Latitudinal distance between native and non-native range centroids was calculated separately for each considered region to which species were introduced and served as a proxy for differences in biomes and community assemblages between native and non-native regions. Again, native ranges and region-specific non-native ranges were based on the WCVP level 3 polygons and/or the GIFT polygons from the status assignment. Last, we considered region-specific introduction history as residence time that was extracted from the database on first occurrence records by Seebens (60). The residence time was calculated as years since the first occurrence record in a region, with 2023 as the reference year. In cases where approximations instead of specific years were provided, we transformed these to numeric values: “early 20^th^ century” to 1925, “mid-20th century” to 1950, and “late 20^th^ century” to 1975.

### Niche change analyses

For each species, we quantified climatic niches and assessed niche changes between each pair of native range and non-native region (hereafter referred to as niche pair). We only considered niche pairs with at least 20 occurrence points within analogue climatic conditions in the native and the non-native niches, respectively. A species was then only included in the overall analysis if we could run the niche comparison between the species’ native range and the Pacific Island region as well as at least one other region. The final analysis covered 316 species and a total of 1566 niche pairs. Of the 316 species that have been introduced to the Pacific Islands, 212 were also introduced to Africa, 173 to temperate Asia, 148 to tropical Asia, 217 to Australasia, 82 to Europe, 199 to North America, and 219 to South America.

Niche changes were quantified using the R package *ecospat* (61), following the methodology of Broennimann et al. (27). Our analysis consisted of three steps, performed for each niche pair. First, the native and non-native realized climatic niches were determined by calculating the occurrence density, and the available environmental space (background data) along the first two axes of a principal component analysis (PCA) over all 19 bioclimatic variables using kernel density estimators. Next, we calculated niche overlap using Schoener’s *D* metric (62) and performed similarity tests (with n = 1200 iterations). The similarity test was used to specifically test for niche conservatism (significantly higher niche overlap than expected by chance, p < 0.05) or niche switching (significantly lower overlap than expected by chance, p < 0.05) between the native and non-native niches. Niche switching is otherwise referred to as niche shift in the literature (see 13), however, we consider the term *shift* to be misleading as to the magnitude of niche changes and thus prefer the term *switching* when speaking of significant deviations from the native niche. We tested these assumptions separately. Lastly, we calculated the niche change in terms of stability, unfilling and expansion. These were standardized to sum up to one, to enable us to assess the relative contribution of each metric to the total analogue niche space. For completeness, we additionally quantified niche abandonment and pioneering in non-analogue climates.

### Trait analyses

We assessed the effect of ecological traits, biogeographic attributes and residence time on region-specific niche changes using multiple phylogenetic regressions. Due to data gaps, only 165 out of the 316 species were included in these analyses, with different numbers of species considered in each region: Africa (n = 124), temp. Asia (n = 95), trop. Asia (n = 78), Australasia (n = 124), Europe (n = 56), N. America (n = 110), Pacific Islands (n = 143), S. America (n = 41). In total, the trait analyses considered 770 of the 1566 quantified niche pairs. All models were estimated using the R package *phylolm* (63), to account for potential biases due to phylogenetic relatedness between the species and their associated traits (64, 65). Phylogenetic data were extracted from the PaciFLora dataset and built on the seed plant phylogeny by Smith and Brown (66). We fitted separate models for each niche change metric and region. In each case, the most parsimonious model was selected using stepwise, Akaike information criterion (AIC)-based variable selection. To determine variable importance, each variable was randomly permutated (n = 99) at a time and the drop in explained deviance between the model with and without permutation was quantified. Variable importance across predictors was standardized to sum up to 100%. We additionally fitted univariate models with higher species sample sizes for each region, to account for the robustness of the signals detected in the most parsimonious models against missing species. We also tested the inclusion of an interaction term between residence time and ecological traits in preliminary analyses. As the interaction terms did not significantly increase explained variance, we did not consider them in the final trait analyses.

## Author Contributions

Anna Rönnfeldt and Damaris Zurell conceived the ideas and designed methodology; Anna Rönnfeldt, Valén Holle and Katrin Schifferle prepared the data; Anna Rönnfeldt analyzed the data; Anna Rönnfeldt led the writing of the manuscript. All authors contributed critically to the drafts and gave final approval for publication.

## Acknowledgements

We acknowledge support from the German Science Foundation DFG [grant no. ZU 361/3-1].

