## Supplementary materials for "Climatic niche conservatism in non-native plants depends on introduction history and biogeographic context"

### Supplementary Figures

##### Map of the study regions (Figure S1)

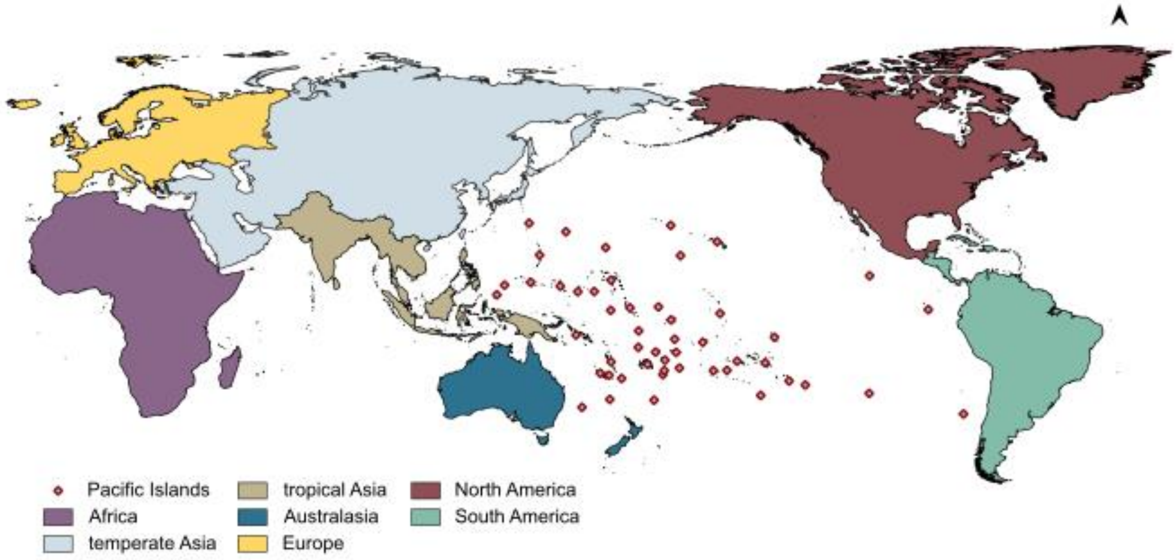

**Figure S1. Map of the eight study regions.** The regions correspond to level 1 of the world geographic scheme of recording plant distributions (1), except for the Pacific Islands which are based on Wohlwend et al. (2) and consist of a subset of 50 island groups.

#### Species flow from native main climate zones to non-native regions (Figure S2)

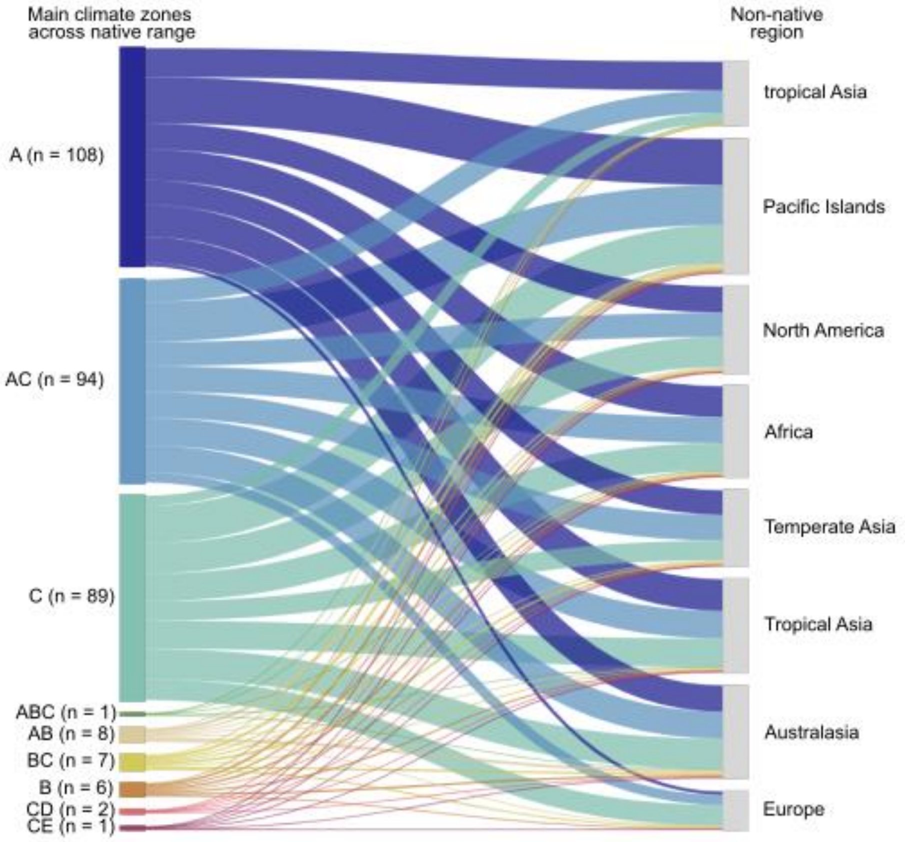

**Figure S2 – Species flows from native main climate zones to non-native regions.** The climate zones are based on the main climate zones of the Köppen-Geiger climate classification *(3)*: A – tropical, B – arid, C – temperate, D – continental, E – polar. A climate zone was considered to be among the main climate zones for a species if 30 % of the species' native occurrences lie within that zone. The number of species associated with the respective climate zones is given in the brackets behind the labels.

**Similarity test outcomes showing the regional split in percentage (Figure S3)**

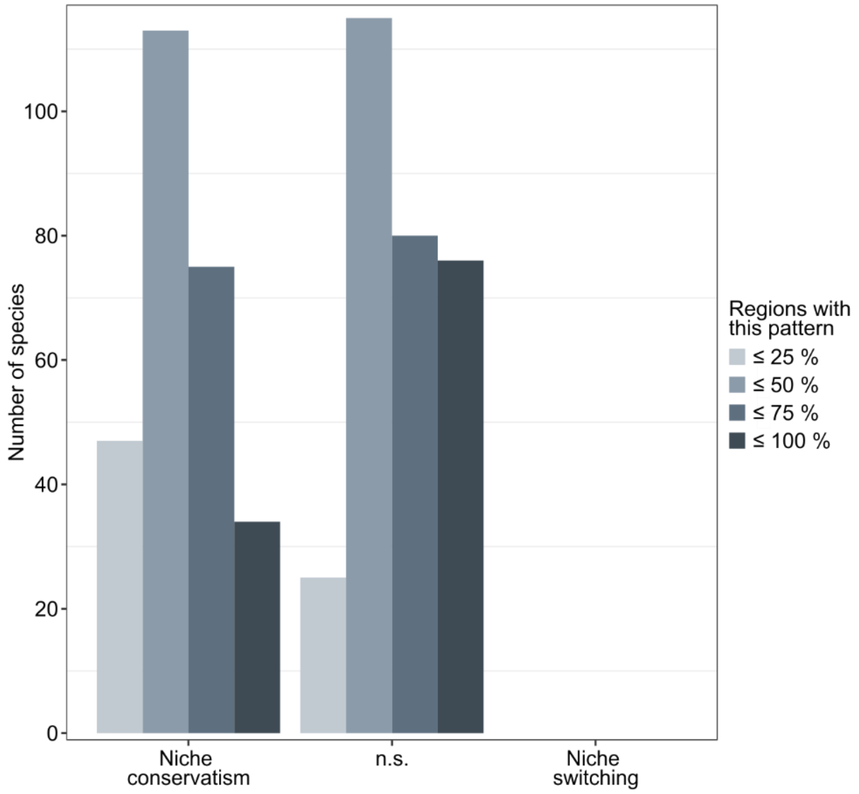

**Figure S3 - Number of species showing climatic niche conservatism or switching.** Significant niche conservatism and switching were determined with similarity tests (n = 1200 iterations). The colored bars indicate the number of species which consistently showed the respective outcomes in x percentage of the regions they have been introduced to.

#### Niche dynamics across analogue and non-analogue niche space (Figure S4)

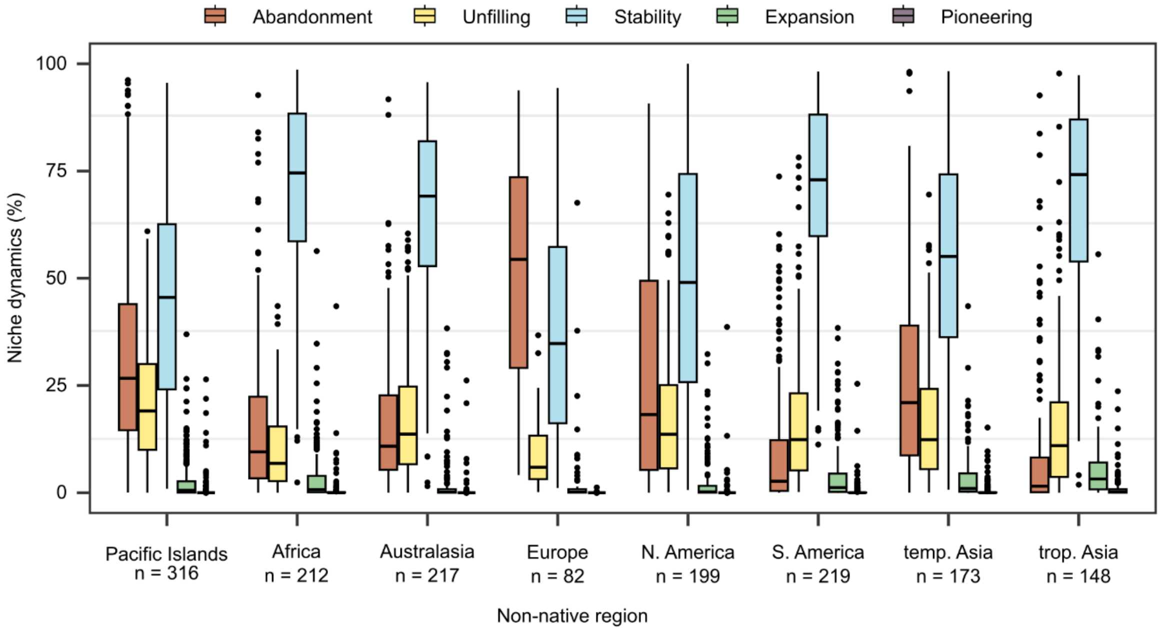

**Figure S4 – Niche dynamics in the non-native ranges.** The sample size under each region name indicates the number of species that have been introduced to that region.

#### Mean trait values for species from different climate zones (Figure S5)

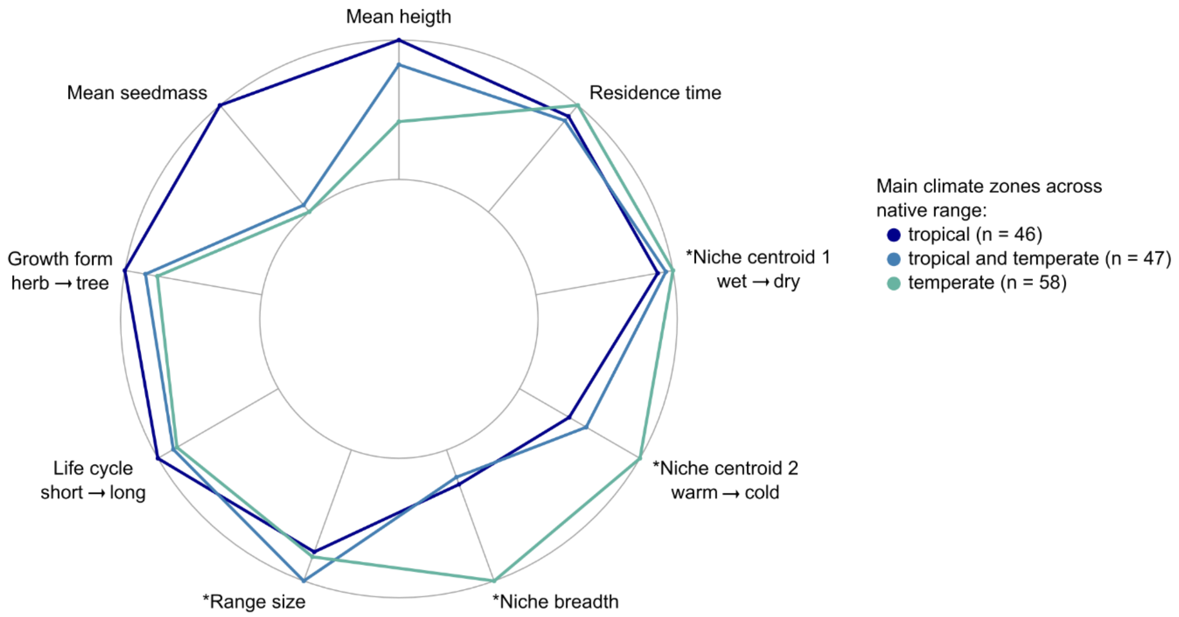

**Figure S5 – Mean trait values for species native to different climate zones.** Traits labelled with an asterisk refer to biogeographic traits estimated for the native range or native niche of the species

#### Trait analysis: results for abandonment and pioneering (Figure S6)

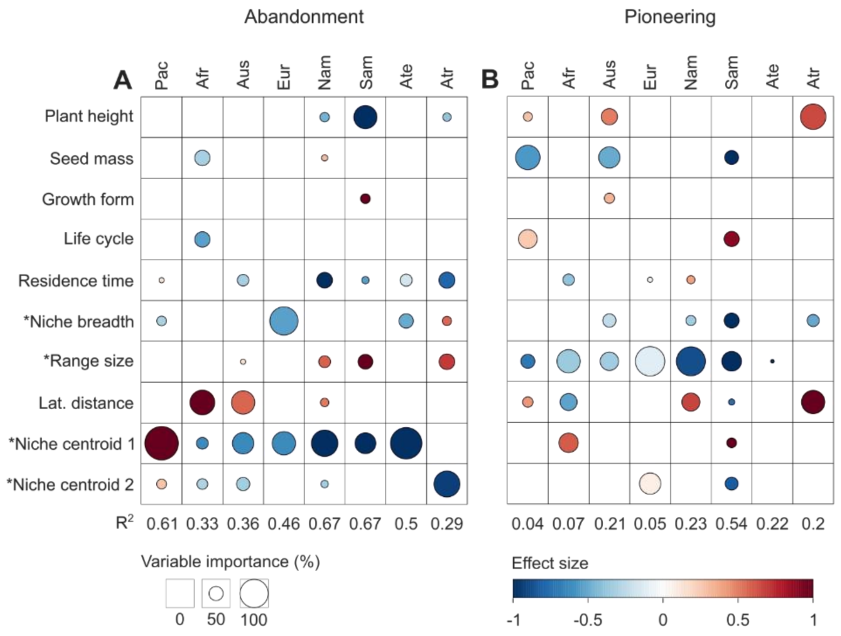

**Figure S6 – Trait effects on the non-analogue niche change metrics in non-native plants.** For each region, we ran AIC-based stepwise phylogenetic regression models with (A) niche abandonment and (B) niche pioneering as responses. The final, parsimonious models per region are shown in each column, with the total explained variance ($R^2$) of the models shown below the columns. Circle sizes indicate the variable importance (%) of single traits within the multiple regression models, and the effect size shows whether the respective niche change metrics will increase (red) or decrease (blue) as the trait values increase. Traits labelled with an asterisk refer to biogeographic traits estimated for the native range or native niche of the species. Niche centroids refer to the relative position along climatic gradient 1 (from warm to cold) and 2 (from wet to dry). The species sample size varied between regions: Africa (Afr, n = 124), temperate Asia (Ate, n = 95), tropical Asia (Atr, n = 78), Australasia (Aus, n = 124), Europe (Eur, n = 56), North America (Nam, n = 110), Pacific Islands (Pac, n = 143), South America (Sam, n = 41).

#### Trait analysis: full models for all niche metrics (Figure S7)

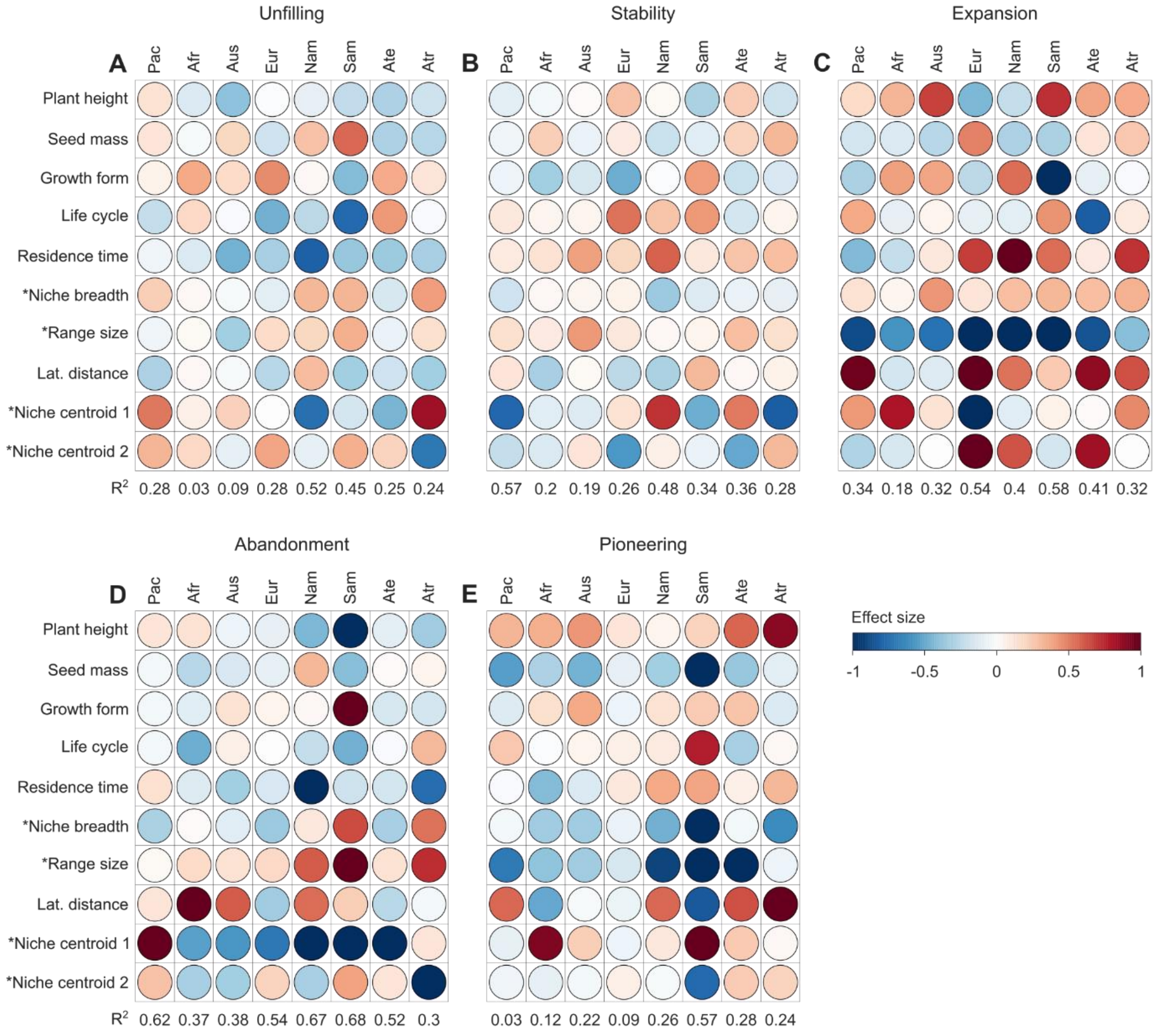

**Figure S7 – Trait effects on the non-analogue niche change metrics in non-native plants.** For each region, we ran phylogenetic regression models with (A) niche unfilling, (B) niche stability, (C) niche expansion, (D) niche abandonment and (E) niche pioneering as responses. Each column shows the full regional model, with the total explained variance ($R^2$) of the models shown below the columns. The effect size shows whether the respective niche change metrics will increase (red) or decrease (blue) as the trait values increase. Traits labelled with an asterisk refer to biogeographic traits estimated for the native range or native niche of the species. Niche centroids refer to the relative position along climatic gradient 1 (from warm to cold) and 2 (from wet to dry). The species sample size varied between regions: Africa (Afr, n = 124), temperate Asia (Ate, n = 95), tropical Asia (Atr, n = 78), Australasia (Aus, n = 124), Europe (Eur, n = 56), North America (Nam, n = 110), Pacific Islands (Pac, n = 143), South America (Sam, n = 41).

#### Trait analysis: univariate models for all niche metrics (Figure S8)

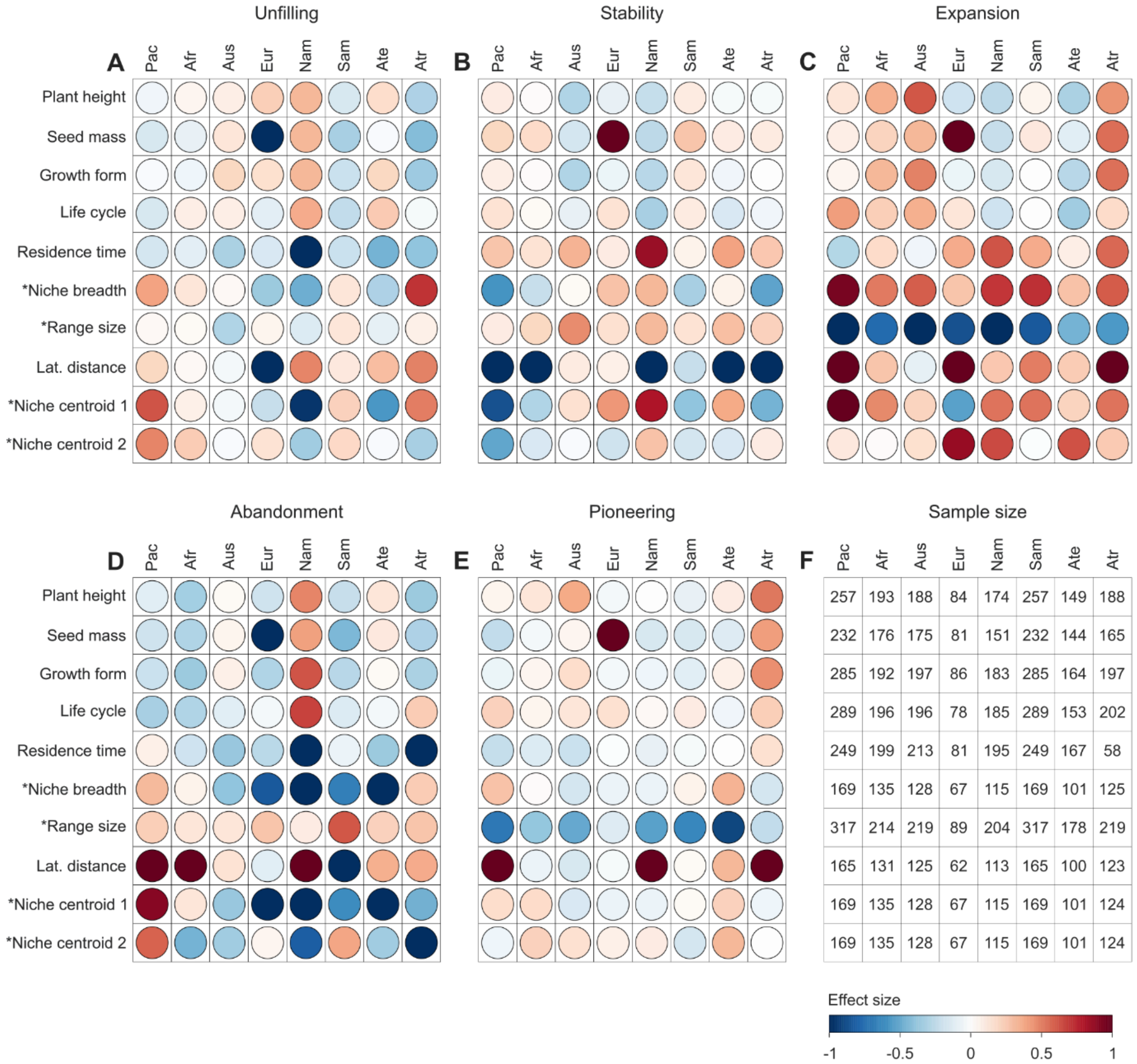

**Figure S8 – Trait effects on the non-analogue niche change metrics in non-native plants.** For each region, we ran univariate phylogenetic regression models with (A) niche unfilling, (B) niche stability, (C) niche expansion, (D) niche abandonment and (E) niche pioneering as responses. The circle color in each cell indicates the effect size for the predictor corresponding to that row and the region corresponding to that column: The effect size shows whether the respective niche change metrics will increase (red) or decrease (blue) as the trait values increases in that region. Traits labelled with an asterisk refer to biogeographic traits estimated for the native range or native niche of the species. Niche centroids refer to the relative position along climatic gradient 1 (from warm to cold) and 2 (from wet to dry). Note that sample size varies across species and trait combinations (F).

### Supplementary Tables

#### ANOVA table for the comparison of the regional niche dynamics (Table S1)

Table 2S – Mean values and ANOVA table for niche unfilling, niche stability, and niche expansion.

| Niche change metric | Region | Mean | ANOVA Estimate | ANOVA Std. Error | ANOVA p-value |
|---|---|---|---|---|---|
| Unfilling | Pacific Islands (Intercept) | 0.34 | -0.65 | 0.12 | ≤ 0.001 *** |
| | Africa | 0.12 | -1.95 | 0.24 | ≤ 0.001 *** |
| | temperate Asia | 0.25 | -1.12 | 0.21 | ≤ 0.05 * |
| | tropical Asia | 0.19 | -1.44 | 0.24 | ≤ 0.001 *** |
| | Australasia | 0.22 | -1.26 | 0.20 | ≤ 0.01 ** |
| | Europe | 0.22 | -1.26 | 0.29 | ≤ 0.05 * |
| | North America | 0.42 | -0.98 | 0.20 | ≤ 0.1 |
| | South America | 0.19 | -1.45 | 0.21 | ≤ 0.001 *** |
| Stability | Pacific Islands (Intercept) | 0.61 | 0.45 | 0.12 | ≤ 0.001 *** |
| | Africa | 0.84 | 1.63 | 0.22 | ≤ 0.001 *** |
| | temperate Asia | 0.71 | 0.88 | 0.20 | ≤ 0.05 * |
| | tropical Asia | 0.75 | 1.08 | 0.22 | ≤ 0.01 ** |
| | Australasia | 0.76 | 1.14 | 0.20 | ≤ 0.001 *** |
| | Europe | 0.74 | 1.02 | 0.28 | ≤ 0.05 * |
| | North America | 0.88 | 0.86 | 0.19 | ≤ 0.05 * |
| | South America | 0.77 | 1.20 | 0.20 | ≤ 0.001 *** |
| Expansion | Pacific Islands (Intercept) | 0.05 | -3.04 | 0.27 | ≤ 0.001 *** |
| | Africa | 0.04 | -3.21 | 0.45 | > 0.1 |
| | temperate Asia | 0.05 | -3.00 | 0.45 | > 0.1 |
| | tropical Asia | 0.06 | -2.74 | 0.44 | > 0.1 |
| | Australasia | 0.02 | -3.77 | 0.53 | > 0.1 |
| | Europe | 0.04 | -3.09 | 0.60 | > 0.1 |
| | North America | 0.03 | -3.66 | 0.53 | > 0.1 |
| | South America | 0.04 | -3.15 | 0.43 | > 0.1 |
